# Using the modified male osteoporosis self-assessment tool for Taiwan to predict osteoporosis onset – a sub-study of the Taiwan osteoporosis survey

**DOI:** 10.1101/479303

**Authors:** Dung-Huan Liu, Tien-Tsai Cheng, Jia-Feng Chen, Shan-Fu Yu, Wen-Chan Chiu, Chung-Yuan Hsu, Ying-Chou Chen

## Abstract

**Purpose:** To develop a risk index by item reduction from multiple variable regression, which can identify male Taiwanese patients at risk of developing osteoporosis.

**Methods:** To develop the model, a risk index was identified by item reduction from multivariate regression analysis. Using receiver operating characteristic (ROC) curve analysis and their sensitivity/specificity, MOSTAi was validated in a separate cohort of Taiwanese men and its performance with compared with the National Osteoporosis Foundation recommendations (NOF 2013).

**Results:** Between 2008 and 2011 a total of 4,323 males were enrolled for bone mineral density (BMD) measurements. Univariate analysis identified four major risk factors for osteoporosis, including age, body weight (BW), previous fracture and body height. The ROC analysis showed the area under the curve (AUC) for the model based on the three-variable, two-variable (age and BW), and one-variable models (BW), was 0.701 (p<0.001, 95% confidence interval [CI] 0.658-0.744), 0.700 (p<0.001, 95% CI, 0.656-0.742), and 0.690 (p<0.001, 95% CI, 0.646-0.734), respectively. Using the optimal cutoff value (−2) for the OSTA, the sensitivity, specificity, positive predictive value (PPV) and negative predictive value (NPV) in the validation cohort were 64.0%, 65.7%, 26.9% and 90.2%, respectively. The ROC curves for predicting osteoporosis by MOSTAi, OSTA and NOF 2013 and the AUC for MOSTAi, OSTA and NOF 2013 was 0.706 (p<0.001, 95% CI: 0.664-0.748) and 0.697(p<0.001, 95% CI: 0.657-0.738), respectively.

**Conclusion:** The results showed that MOSTAi could be a more precise model than OSTA and NOF 2013, for identifying men in Taiwan with osteoporosis who require referrals for DXA scans. It was demonstrated that MOSTAi is a simple tool with fair sensitivity/specificity and PPV, and high NPV. MOSTAi could also be a more accurate model than OSTA for identifying men in Taiwan at risk of osteoporosis. In comparison with NOF 2013, MOSTAi is a more accurate and simpler tool for the referral of Taiwanese men for DXA scans.

## Introduction

As the world population is aging, osteoporosis in males is becoming a global health concern. Aging men lose around 1% bone mineral density (BMD) per year [1,2], and 20% of men aged over 50 will develop an osteoporotic related fracture during their lifetime [3,4]. In Taiwan the percentage of the population aged over 50 years is predicted to grow from 32% in 2013 (7.5 million) to 57% in 2050 (11.9 million), and the life expectancy of 80 years in 2013 is predicted to increase to 83 years by 2050 [5]. The annual hip fracture rate for Taiwanese men was higher in Asian regions [6]. In addition, 22% of Taiwanese men with a hip fracture died within a year of their injury, which is notably higher when compared with women who suffer hip fractures. Therefore, the identification of men who have osteoporosis is mandatory for the prevention of osteoporotic related fractures.

According to the World Health Organization (WHO) classification criteria, dual-energy X-ray absorptiometry (DXA) is the gold standard diagnostic test for osteoporosis (T-score, −2.5). Due to health care reimbursement restrictions and the limited availability of DXA machines in some rural areas, regular DXA scans for BMD are not available to all of the Taiwanese population. To improve the targeted, cost-effective use of DXA, several tools have been developed [7–15], including the osteoporosis self-assessment tool (OSTA) [7] and National Osteoporosis Foundation (NOF 2013) [15].

Since 1993 several studies have demonstrated that aging and low BW are associated with osteoporosis fractures [16–19]. In 2001 Koh *at el*. proposed OSTA, which is an easy method of identifying those at risk of osteoporosis using BW and age [7]. NOF 2013 has also been used to identify individuals who are at high risk of developing osteoporosis and should arrange a DXA examination. At present OSTA has been validated in men from a variety of races [20–27], however it has not yet been directly evaluated in Taiwanese men. In addition, OSTA has not yet been directly compared with NOF 2013 for its effectiveness in men. Therefore, the present study developed a risk index named MOSTAi and validated it by using a separate cohort and by comparing it with the NOF 2013 recommendations.

## Materials and methods

### Participants

Between 2008 and 2011 the Taiwan osteoporosis association (TOA) performed a whole island circuit program for the assessment of BMD. This program included a trained nurse, a bus installed with a DXA machine (Explorer; Hologic Inc., Waltham, MA, USA) International Society of Clinical Densitometry (ISCD) and a certified radiology technician. The bus travelled around districts on request and took BMD measurements using the DXA machine. A total of 104 sites were enrolled across Taiwan.

For inclusion with the study patients were required to fulfill the following inclusion criteria: male, aged ≥50 years, with a hip anatomy suitable for assessment via a DXA scan and provided informed consent. If the patient had both hips replaced or previously fractured, or could not access the DXA machine they were excluded from the study.

### Data collection and measurements

The trained nurse interviewed each participant and asked them to complete the list of questions in the Fracture Risk Assessment (FRAX) tool. Risk factors including BW, body height (BH), glucocorticoid use, previous fragility fracture, rheumatoid arthritis (RA), secondary osteoporosis, gender, age, parent with a fractured hip, smoking and alcohol consumption were assessed.

On the bus the DXA machine was used to take BMD measurements of the hip regions and lumbar spine in all patients. The reference for those aged 20-29 years was used for BMS values [28], which is the recommended reference database for local Taiwanese men.

The subjects were assigned to one of two groups: (1) the non-osteoporotic risk group or (2) those at risk of developing osteoporosis in need of BMD measurement. Patients were assigned to the at risk group if they met one of the following criteria: male aged 50-69 with a previous fracture, men aged ≥70 years with a low BWI (BMI <18.5 kg/m^2^), the use of high risk medication associated with low bone mass or bone loss (e.g. glucocorticoid steroids at daily doses ≥5 mg prednisone or equivalent) for ≥3 months, or disease associated with bone loss (e.g. RA). NOF 2013 was then used to evaluate the risk of future osteoporosis.

### Statistical analysis

The t-test was used for continuous variables and the chi-square test was used for categorical variables. The FRAX tool was used to assess possible risk factors in the model development. Firstly, univariate analyses were performed and the statistically significant risk factors were identified (p<0.01) and then included in the multiple variable regression model. The next step was to develop a simple task by major risk factors via multiple variable regression analysis and the item reduction method.

ROC curve analysis was performed to assess the ability of MOSTAi to distinguish between subjects with and without osteoporosis. The performance of MOSTAi was compared with OSTA and NOF 2013 using the AUC. All analyses were performed using SPSS statistics. A p-value <0.01 was considered to indicate a statistically significant difference.

## Results

A total of 18,992 participants, including 4,323 males (22.8%) and 14,669 females (77.2%) were enrolled in the Taiwan Osteoporosis Survey (TOPS). Patients whose BMD data was unavailable, those whose FRAX-based question data was incomplete or missing, all females, and males aged <50 years were excluded from the analysis. The distribution of participants is shown in Figure 1. Overall 2,290 men were evaluated and randomly assigned to either the development (n=1145) or validation (n=1145) cohorts.

**Fig 1.**
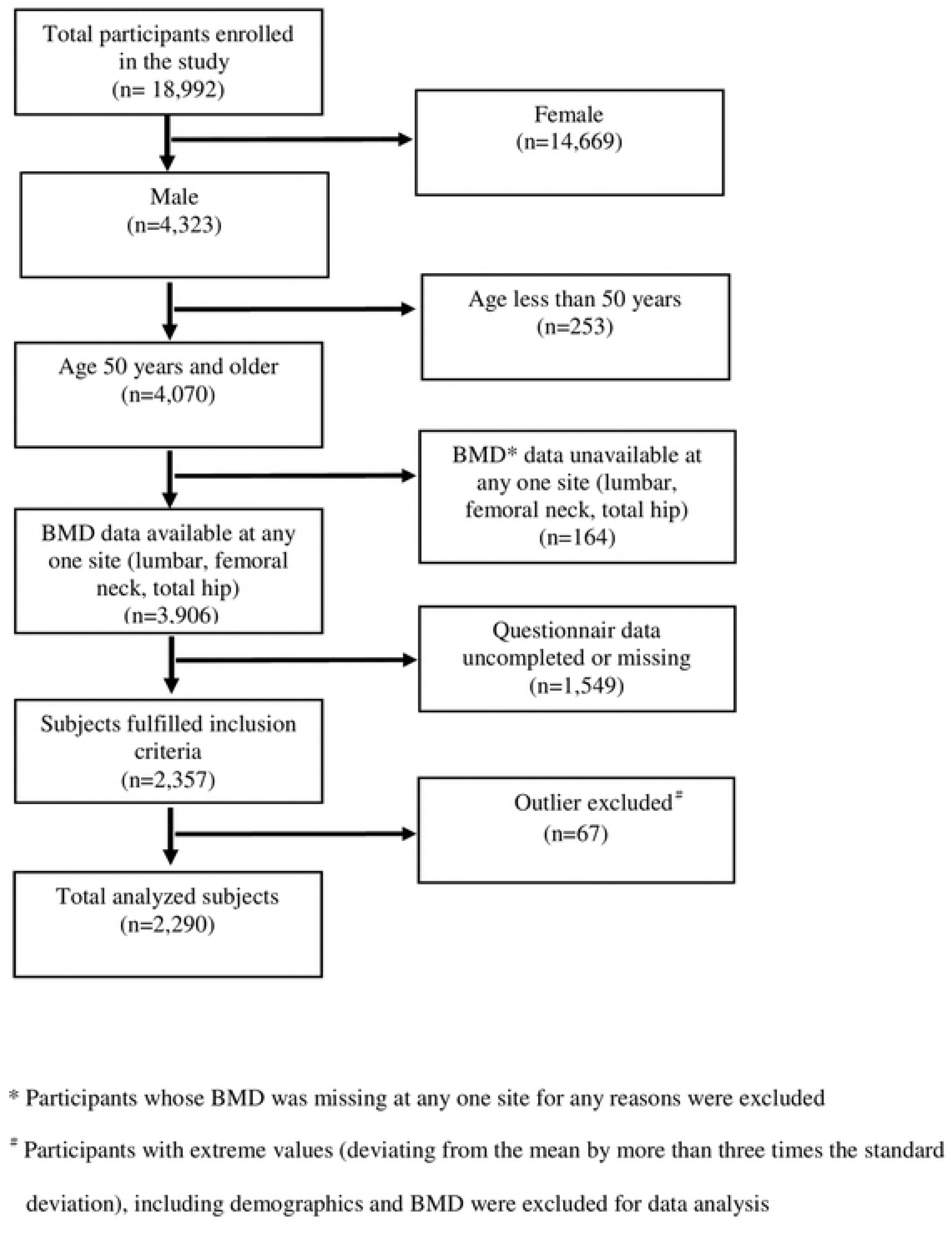
The inclusion of participants in the study.

Characteristics of the development and validation cohorts are shown in Table 1. Univariate analysis was performed and four major risk factors for osteoporosis were identified, including age, BW, previous fracture and body height. The index weights for the three factors as determined by multivariate regression were ultimately used to score the MOSTAi index value for each subject. The regression coefficient for both multivariate and univariate analysis in the development cohort are shown in Table 2.

**Table 1.** Demographic characteristics of all participants.

| Characteristics | Total number<br>n=2290 (100%) | Development<br>Cohort<br>N=1145 (50%) | Validation<br>Cohort<br>N=1145 (50%) | P-value |
| --- | --- | --- | --- | --- |
| Age (years) | 69.6±9.6 | 69.9±9.5 | 69.4±9.6 | 0.294 |
| Height (cm) | 165.4±6.1 | 165.5±6.0 | 165.2±6.1 | 0.323 |
| Weight (kg) | 65.9±9.8 | 65.9±10.1 | 65.8±9.6 | 0.727 |
| Body mass index<br>(kg/m<sup>2</sup>) | 24.1±3.2 | 24.0±3.3 | 24.1±3.1 | 0.868 |
| BMD (g/cm<sup>2</sup>)<br>(T-score) |  |  |  |  |
| Lumbar spine (g/cm<sup>2</sup>)<br>(T-score)* | 1.01±0.15<br>-0.7±1.4 | 1.02±0.15<br>-0.7±1.4 | 1.01±0.15<br>-0.7±1.4 | 0.950 |
| Femoral neck (g/cm<sup>2</sup>)<br>(T-score)* | 0.76±0.13<br>-1.2±1.0 | 0.76±0.13<br>-1.2±1.0 | 0.76±0.13<br>-1.3±1.0 | 0.777 |
| Total hip (g/cm<sup>2</sup>)<br>(T-score)* | 0.93±0.15<br>-0.7±1.0 | 0.93±0.15<br>-0.7±1.0 | 0.93±0.15<br>-0.7±1.0 | 0.988 |
| **Other risk factors\* in FRAX<sup>®</sup>, n/ N@** |  |  |  |  |
| Parent Fractured Hip<br>(%) | 228/2290(10.0) | 110/1145(9.6) | 118/1145(10.3) | 0.577 |
| Previous Fracture (%) | 117/2290(5.1) | 55/1145(4.8) | 62/1145(5.4) | 0.507 |
| Glucocorticoids (%) | 107/2290(4.7) | 55/1145(4.8) | 52/1145(4.5) | 0.767 |
| Rheumatoid arthritis<br>(%) | 111/2290(4.8) | 59/1145(5.2) | 52/1145(4.5) | 0.496 |
| Secondary<br>osteoporosis (%) | 150/2290(6.6) | 80/1145(7.0) | 70/1145(6.5) | 0.617 |
| Current smoking (%) | 372/2290(16.2) | 191/1145(16.7) | 181/1145(15.8) | 0.571 |
| Alcohol 3 or more units/day (%) | 115/2290(5.0) | 52/1145(4.5) | 63/1145(5.5) | 0.293 |

\* Definition same as those of FRAX
@ n=answered yes in the questionnaire, N= total number who answered the questionnaire .

**Table 2.** Regression coefficients for the univariate and multivariable analysis in the development cohort.

| | $\beta$ | SE* | p | $\beta$ | SE* | p | Index weight |
| --- | --- | --- | --- | --- | --- | --- | --- |
| Age (vs. 10 years younger) | -0.172 | 0.030 | <0.001 | -0.092 | 0.029 | 0.001 | -1 |
| Body weight (vs. 10 kgs lighter) | 0.358 | 0.027 | <0.001 | 0.325 | 0.030 | <0.001 | 3 |
| Previous fracture (vs. no) | -0.106 | 0.134 | <0.001 | -0.086 | 0.125 | 0.002 | -1 |
| Body height (vs. 10 cm shorter ) | 0.194 | 0.047 | <0.001 | 0.022 | 0.050 | 0.471 |  |
| Parent hip fracture (vs. no) | 0.016 | 0.098 | 0.579 | - | - | - |  |
| Glucocorticoids (vs. no) | -0.010 | 0.135 | 0.736 |  |  |  |  |
| Rheumatoid arthritis (vs. no) | -0.016 | 0.130 | 0.592 | - | - | - |  |
| Secondary osteoporosis (vs.no) | -0.021 | 0.113 | 0.482 | - | - | - |  |
| Smoking (vs. no) | -0.038 | 0.077 | 0.202 |  |  |  |  |
| Alcohol (vs. no) | 0.013 | 0.138 | 0.661 | - | - | - |  |

\*SE: standard error

The number of men with a T-score ≤-2.5 at the femoral neck, lumbar spine or the total hip, or a combination of two or all three sites in the development and validation cohorts are shown in Table 3. A total of 373 individuals (16.3%) were assessed as having osteoporosis (T-score ≤ −2.5 at the femoral neck, lumbar spine, or total hip) according to the WHO definition. These compatible to the total number of subjects with T-scores at −2.5 or below at either site in the validation and development cohorts.

**Table 3.**
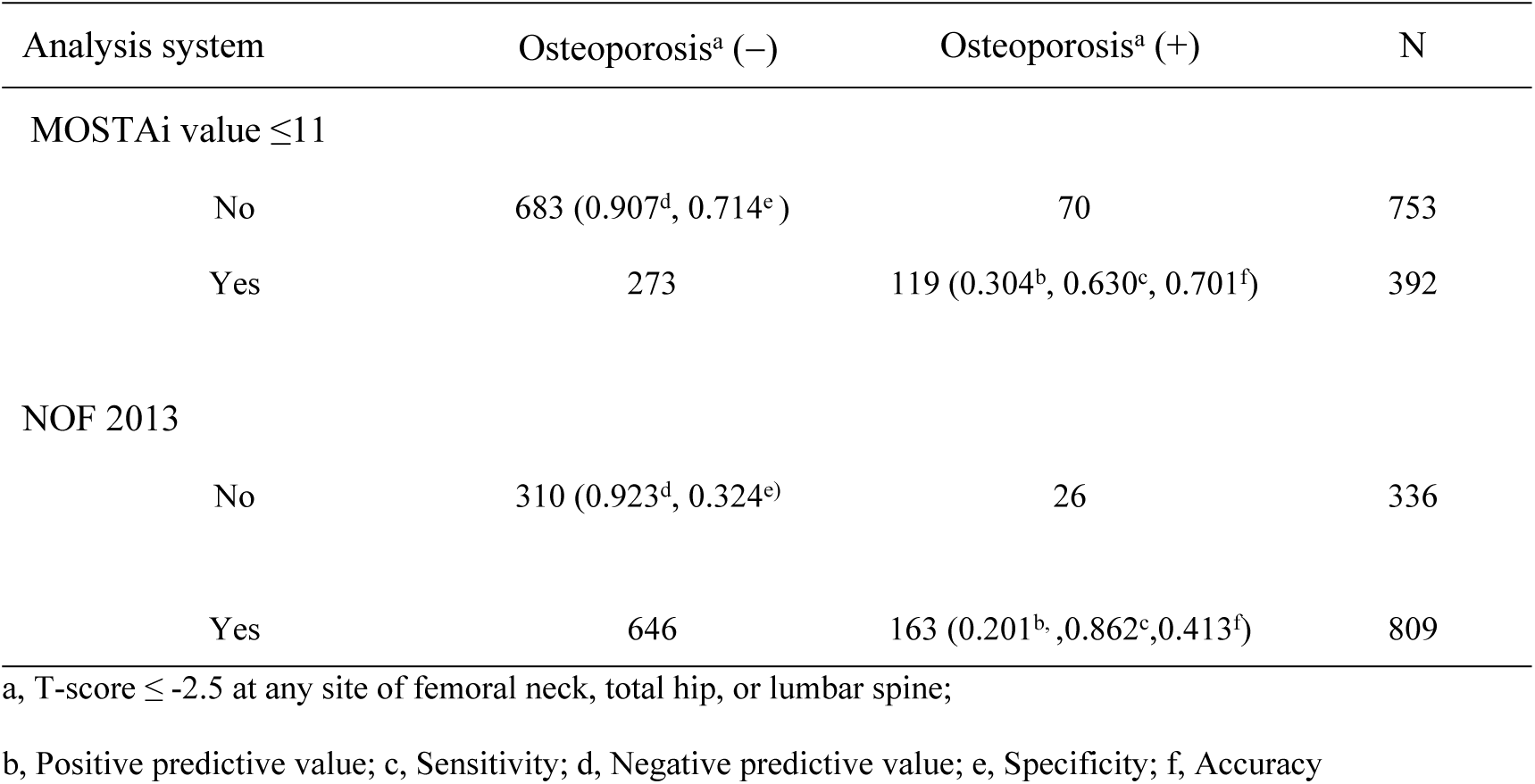
Comparison between the MOSTAi index and NOF 2013 in the validation cohort.

The ROC curve of the development cohort is illustrated in Figure 2 (left side). The final variables were age, BW, and previous fracture. The AUC for the three variables, two variables (age and BW), and one variable (BW) was 0.701 (p<0.001, 95% CI, 0.658-0.744), 0.700 (p<0.001, 95% CI 0.656-0.742), and 0.690 (p<0.001, 95% CI, 0.646-0.734), respectively. A diagnostic tool is poor and considered unacceptable if its AUC is under 0.7 [29]. Due to this the model based on BW only was excluded. By item reduction, performance of the model containing only age and BW was the same as with all three variables. Therefore, the index MOSTAi value for BW and age could be determined by adding +3 units per 10 kg increase in BW to the referent age (50-years-old) and BW (50 kg) and −1 unit per 10 years increase in age. We first count age by weight of −1, and then multiply the last digit, before adding to the index. A similar process was used for BW. The formula for the MOSTAi index value was as follows: 0.3 × (BW in kg) − 0.1 × (age in years). The MOSTAi index values of the 1,145 participants in the development cohort were calculated. The mean, median, standard deviation (SD), and range of the MOSTAi index were 12.8, 13, 3.4, and 4-23, respectively.

**Fig 2.**
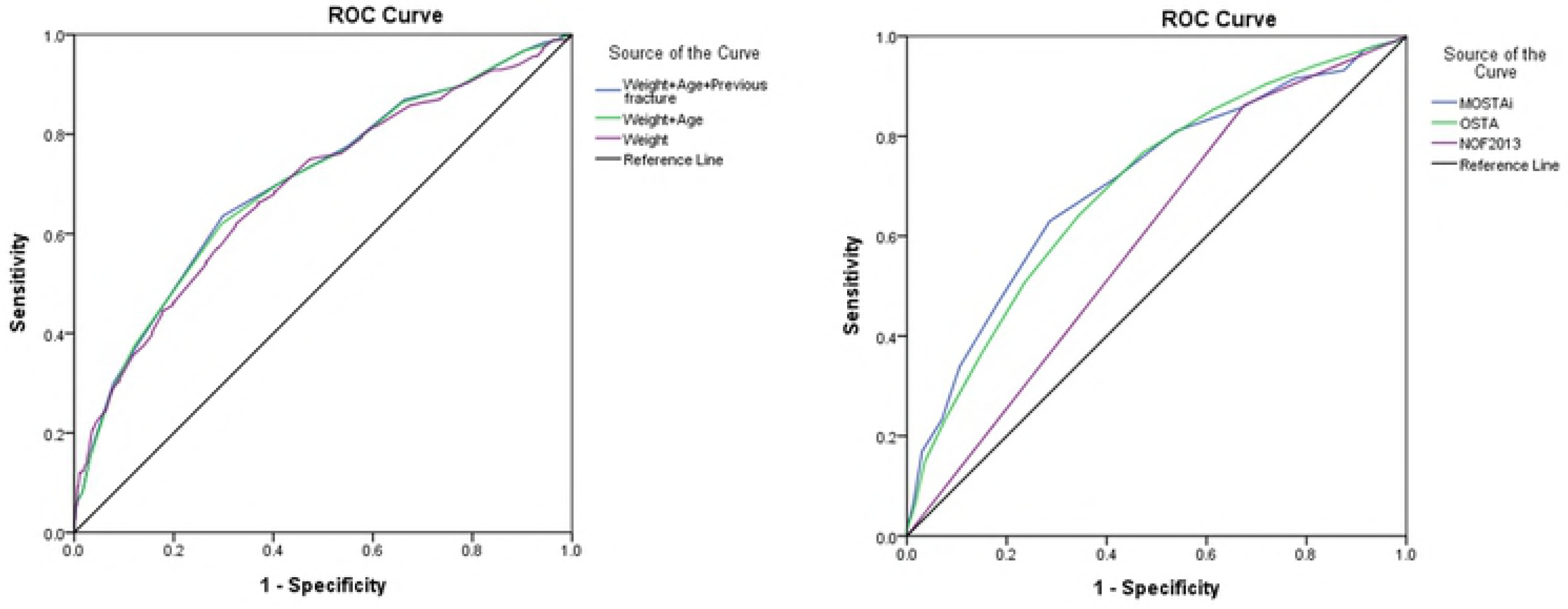
The ROC curves for predicting osteoporosis by MOSTAi, OSTA, and NOF 2013 (right side). The AUC for MOSTAi, OSTA and NOF 2013 were 0.706 (95% CI: 0.664-0.748, p<0.001), 0.697(95% CI: 0.657-0.738, p<0.001) and 0.593(95% CI: 0.552-0.634, P<0.001), respectively. CI, confidence interval; ROC, receiver operating characteristic; AUC, area under the curve; OSTA, osteoporosis self-assessment tool; NOF, National Osteoporosis Foundation; MOSTAi, modified male osteoporosis self-assessment tool for Taiwan.

After estimation of the ROC curve, 11 was selected as the appropriate cutoff for the MOSTAi index to identify subjects at high risk of developing osteoporosis in the development cohort. The sensitivity, specificity, positive predictive value (PPV) and negative predictive value (NPV) of MOSTAi in the development sample (n=1,145) were 61.96%, 70.45%, 28.64% and 90.63%, respectively. A comparison between MOSTAi and NOF 2013 in the validation cohort is shown in Table 3.

Using the optimal cutoff value (−2) for OSTA, the sensitivity, specificity, PPV and NPV in the validation cohort were 64.0%, 65.7%, 26.9% and 90.2%, respectively. The ROC curves for predicting osteoporosis by NOF 2013, MOSTAi and OSTA are shown in Figure 2 (right side). The different AUCs for MOSTAi, OSTA and NOF 2013 were 0.706 (p<0.001, 95% CI: 0.664-0.748), 0.697(p<0.001, 95% CI: 0.657-0.738) and 0.593 (p<0.001, 95% CI: 0.552-0.634), respectively.

Three osteoporosis risk categories were created based on the MOSTAi index. The lowest T-scores at any site and the MOSTAi values for the development samples are shown in Figure 3. The high-risk group included those with a MOSTAi index ≤5, the medium-risk group included those with MOSTAi index values between 5 and 11 (MOSTAi ≤11 and >5), and the low-risk group included those with MOSTAi index values >11.

In the development cohort, 65.2% of patients were classified as low-risk 33.5% were classified as medium-risk and 1.2% were classified as high-risk. The prevalence of osteoporosis was 9.4% (70/747) in the low-risk group, 26.8% (103/384) in the medium-risk group and 78.6% (11/14) in the high-risk group. In the validation cohort, 65.8% of patients were in the low-risk category, 33.1% were in the medium-risk category and 1.1% were in the high-risk category. The estimated prevalence of osteoporosis in the high, medium and low-risk categories were 53.8% (7/13), 29.6% (112/379) and 9.3% (70/753), respectively,.

**Fig 3.**
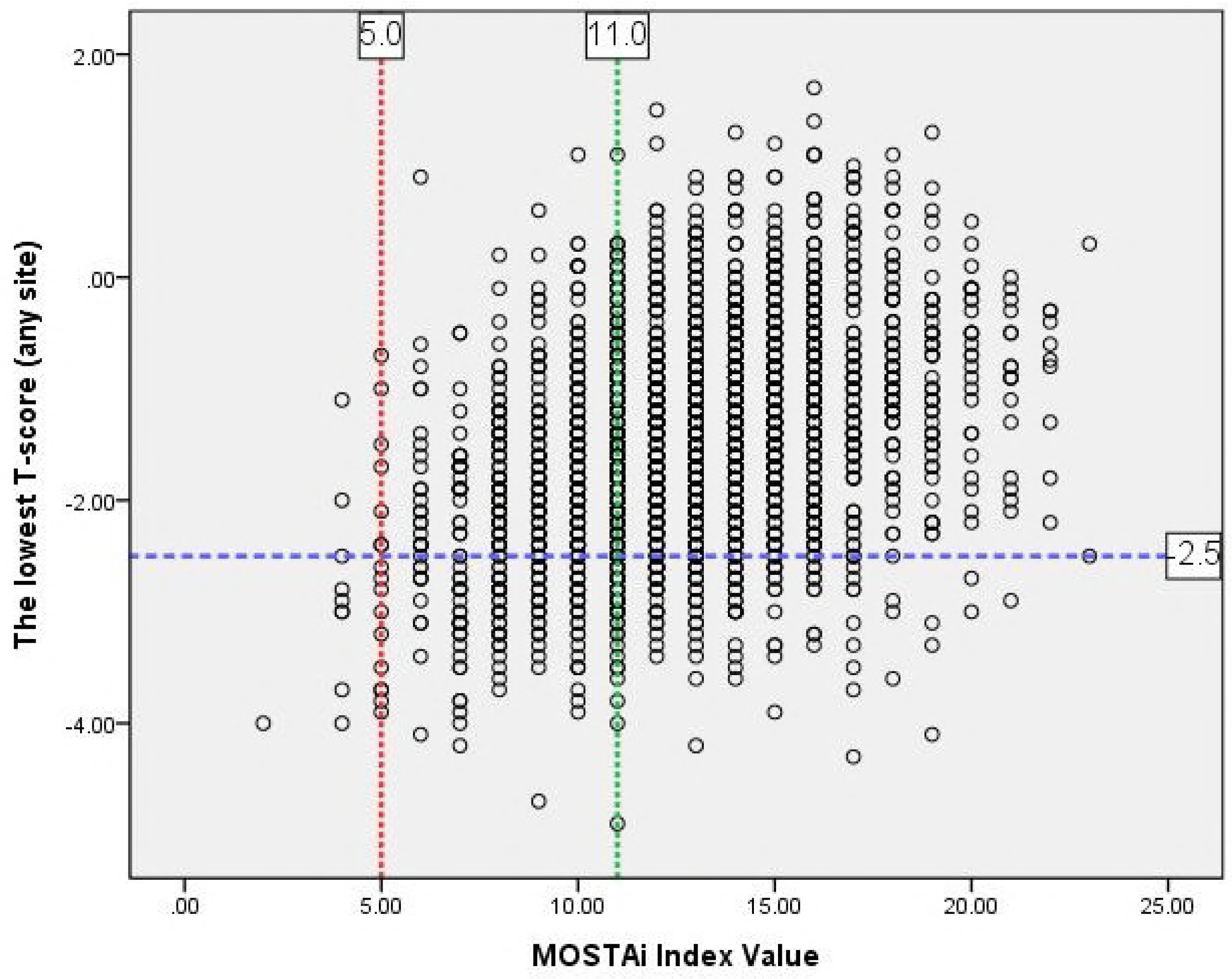
MOSTAi values versus lowest T-scores at any site for the development sample.

## Discussion

OSTA was originally developed using an Asian cohort and later validated in white cohorts [30–32]; it has been effective at identifying postmenopausal women at risk of developing osteoporosis [8]. Recently, the OSTA was further developed to predict osteoporosis in males. As yet, OSTA was validated in multiracial men [20–27,33,34]. Although several studies indicated that OSTA is an effective and simple clinical risk assessment tool for predicting osteoporosis in males [20,21,24,26,27,33,34] Perez-Castrillon *et al.* [35] could not demonstrate that the OSTA index was a useful tool (specificity of 86%, sensitivity of 39%, and a non-significant ROC curve) for identifying osteoporosis in a cohort of 67 Spanish men (mean age, 51 years).

A difference in the accuracy of OSTA for identifying osteoporosis in males compared with females has been reported [26,27]. Richards *et al*. [26] found that the OST index performed better in non-Hispanic whites, and males ≥65 years. Lynn *et al*. [27] observed that the OST index was useful in both populations when using a cutoff ≤ 2 for American Caucasian men and a cutoff of ≤-1 for Hong Kong Chinese men. In addition, a previous study [36] that directly applied the original OSTA model to a cohort of 98 Singaporean men demonstrated that the OSTA performed better using a cutoff value of 0 rather than −1. Therefore, the optimal cutoff for OSTA to predict male osteoporosis may vary in different ethnic groups. As well as ethnicity, the varied performance of OSTA for predicting osteoporosis is males, may be associated with how BMD was measured between investigations [24].

The validation of OSTA in males from a variety of different ethnic groups and MOSTAi in Taiwanese men is summarized in Table 4. In terms of sensitivity/specificity and AUC, the results of OSTA were varied between populations in different studies. In addition, only some of the male OSTA validation studies [20,26,27,34] reported that the AUC was >0.7. Therefore, the application of OSTA for different populations highlights the need to define appropriate cutoffs.

**Table 4.**
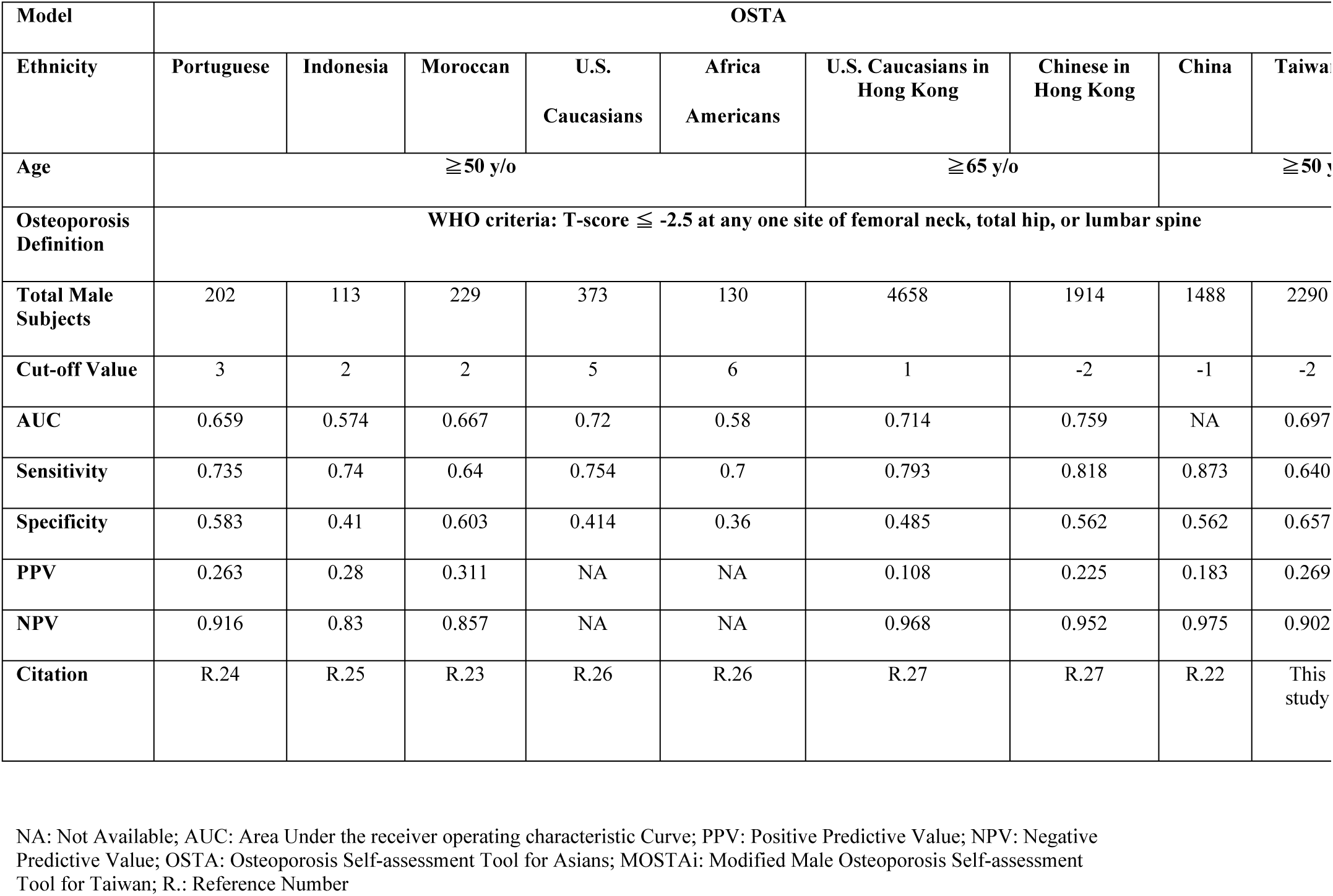
Validation of OSTA in males from multiple races and MOSTAi in Taiwanese men.

In the present study, 373 participants were diagnosed with osteoporosis based on a T-score ≤ −2.5 at either the femoral neck, the lumbar spine or the total hip. The total number of participants with osteoporosis would have been underestimated if measurements were taken at the lumbar spine only (n=127), the femoral neck only (n=117), or at the total hip only (n=7). The effectiveness of OSTA for identifying osteoporosis may depend on the BMD measurement site. Ghazi *et al*. [23] and Saraví *et al*. [37] observed that the appropriate OSTA index cutoff may be based on different skeletal sites. In addition, osteoporosis at one site can predict future fracture risk at other sites [38].

As osteoporosis risk factors and their weighting index may differ depending on the patient’s sex and ethnicity, adjustments may be necessary to make OSTA applicable to men aged ≥50 years old. There is a worldwide trend to commence treatment for osteoporosis by a fracture risk assessment as opposed to BMD alone [25]. The FRAX tool [39] was launched in 1998 and is based on the use of clinical risk factors. Therefore, in the current investigation, the participants were asked to complete a questionnaire, which included the 11 elements in the FRAX tool. As some of the elements in the FRAX tool are also risk factors for osteoporosis, they were incorporated for the development of MOSTAi for men.

In a previous study by the authors [40], some modifications were made to the original OSTA model by including risk factors from FRAX; this led to the development of OSTAi for Taiwanese postmenopausal women, which used the same algorithm as the original OSTA model. In the present study, the algorithm for calculating MOSTAi is similar to the original OSTA formula, except it uses different index weights. The weighting index of age and BW were −1 and ＋ 3 in the MOSTAi model. MOSTAi values of ≤11 yielded a sensitivity of 61.9% and a specificity of 70.45%, and the AUC was 0.700 (95% CI, 0.656-0.742, P<0.001). The MOSTAi index was further validated in another sample of 1,145 Taiwanese men with similar results (sensitivity, 63.0%, specificity of 71.4%, and AUC, 0.706).

In the present study, increased age and a lower BW were independent risk factors for osteoporosis, which was in keeping with findings by Kung *et al*. [34] in a cohort of 420 Chinese men (age ≥50 years). However, the present study found that BW had a higher contribution (index weight = 3) to the MOSTAi index value compared with the investigation by Kung *et al*. The reason why a low BW played a more significant role in the development of osteoporosis in the elderly male Taiwanese cohort compared with the Chinese cohort is unknown and requires further investigation.

The performance of MOSTAi was further compared with OSTA and NOF 2013 in the validation sample of 1,145 Taiwanese men. The optimal cutoff value of −2 for OSTA yielded a sensitivity and specificity of 64.0% and 65.7%, respectively and the AUC was 0.697. In the validation cohort MOSTAi showed acceptable sensitivity/specificity (63.0/71.4%) and a high negative predictive value (90.7%). So as to reduce unnecessary tests and costs for patients, the specificity must be contained within reasonable levels. A high negative predictive value can screen out Taiwanese men with a low risk of osteoporosis, just like male OSTA in other studies. Therefore, MOSTAi could be a more appropriate tool than OSTA for the identification of Taiwanese men at risk of developing osteoporosis. The accuracy (69.6%) and AUC (0.706) of MOSTAi were increased compared with those of NOF 2013 (40.2% and 0.593, respectively). MOSTAi may be an easy to use, more effective screening tool.

The application of three risk categories provides a useful alternative to a single cutoff approach when there is a spectrum of risk, as with osteoporosis [7]. In the present study, 78.6% of the high-risk category (MOSTA index values ≤5) in the developmental group had osteoporosis. It may be appropriate to refer patients who fall within this group for DXA scanning. At the other extreme, because the prevalence of osteoporosis in the low-risk group (9.4% in the development cohort) was low, it would be reasonable to defer BMD measurements for patients within this group. Although the prevalence of osteoporosis in the medium-risk group (26.8% in the development cohort) was not high enough to strongly recommend BMD measurements, a DXA scan is advised for subjects with additional risk factors for fracture, such as presence of a disease or condition associated with bone loss (e.g. RA).

To date, the TOPS is the first countrywide investigation of osteoporosis in Taiwan, which may be used to launch a more reliable diagnostic tool. It should be noted that all measurements were performed by the same DXA machine and the same technician, which lessens inter-modality and inter-operator variation. Furthermore, the risk factors of osteoporotic fracture used in the FRAX tool were included in MOSTAi, whereas they were not used to develop the OSTA for men. The present study did have some limitations. For example, the populations were not randomly selected and the proportion of osteoporosis may be higher compared with the general population. That is because certain patients were referred by a clinician for osteoporosis evaluation, which could result in selection bias. However, the prevalence of osteoporosis (16.3%) in Taiwanese men aged ≥50 years in the present study was close to that of previous survey (17.2%) in elderly Taiwanese men [41]. Although a lower BW and aging can predict future fracture risk [42], whether MOSTAi can predict the future fracture risk for Taiwanese men requires further investigation.

In conclusion, it was demonstrated that MOSTAi is a simple tool with fair sensitivity/specificity and PPV, and high NPV. It may also be a more appropriate model than OSTA for the identification of Taiwanese men at risk of osteoporosis. In comparison with NOF 2013, MOSTAi is a better and simpler tool for the DXA referral of Taiwanese men.

## Author Contributions

Conceptualization: DUL, TTC

Data curation: JFC, SFY, WCC, CYH

Formal analysis: JFC

Funding acquisition: SFY

Investigation: WCC

Methodology: CYH

Project administration: JFC, SFY

Resources: WCC, CYH

Software: CYH

Supervision: JFC

Validation: JFC

Visualization: WCC, CYH

Writing (original draft preparation): DUL, TTC

Writing (review and editing):YCC

## Acknowledgements

The authors would like to thank the TOA for authorizing access to the database, and Merck Sharp & Dohme pharmaceutical company (Taiwan) for providing the mobile DXA machine during the recruitment period. The authors would also like to thank thanked Professor Chan SJ for their help with the in statistical analysis.

## Author contributions

Dung-Huan Liu, and Tien-Tsai Cheng: Conception and design, data analysis and interpretation, manuscript preparation and revision. Ying-Chou Chen, Jia-Feng Chen, Shan-Fu Yu, Wen-Chan Chiu and Chung-Yuan Hsu: Collection and assembly of data. Tien-Tsai Cheng: Administrative and financial support. All authors read and approved the final manuscript.

